# Non-cyclic dinucleotide STING agonists abrogate MPXV infection

**DOI:** 10.64898/2026.06.08.730963

**Authors:** Lucy Eke, Alazne R. Unanue, Alasdair J.M. Hood, Telma M Sancheira Freitas, Antonio Alcami, Carlos Maluquer de Motes, Bruno Hernaez, Rebecca P. Sumner

## Abstract

Mpox has emerged as a global threat to public health following several national and international outbreaks from 2022. Poxviruses deploy multiple strategies to counteract host immune defences including pathways leading to interferon (IFN) production. Here we demonstrate that depleting the viral 2’3’-cGAMP nuclease poxin restores activation of STING and IRF3 during MPXV infection despite the presence of other viral antagonists. We then demonstrate that non-cyclic dinucleotide (non-CDN) STING agonists are resistant to poxin; activate STING and IRF3 during infection; and potently suppress MPXV and orthopoxvirus replication in human primary fibroblasts and differentiated monocytes, where replication is completely abrogated. Mechanistically, non-CDN restriction requires STING, IFNAR and STAT signalling, and induces a unique transcriptional signature over IFNβ, enabling expression of additional cytokines. In vivo, non-CDN activity effectively reduces signs of illness and enhances survival in wild-derived castaneous mice inoculated with the virulent clade I MPXV. Our study reveals poxin as an Achilles’ heel of MPXV, that when bypassed by direct STING agonism, provides a promising novel anti-mpox therapeutic strategy with reduced risk of antiviral resistance.

## Introduction

Orthopoxviruses (OPXV) are a genus of large DNA viruses that replicate exclusively in the cell cytoplasm and infect a variety of rodent species, with some gaining the ability to infect and transmit amongst humans and cause disease. This includes variola virus (VARV), the causative agent of smallpox, and monkeypox virus (MPXV), the causative agent of mpox. Whilst smallpox has been eradicated, mpox cases have increased significantly in recent years, posing an ever-increasing epidemic and pandemic risk (1). A global outbreak of mpox began in May 2022 caused by a clade IIb MPXV strain (lineage B.1) genetically linked to endemic strains identified in West Africa (2–4). This outbreak has resulted in continuous community transmission in multiple non-endemic countries to this day, with >100,000 infections reported. In addition, two large outbreaks are ongoing in the Democratic Republic of the Congo (DRC) and Sierra Leone with spillover events in surrounding countries (5, 6). Whilst the outbreak in Sierra Leone is caused by a clade IIb lineage A virus, the outbreak in the DRC is caused by a strain belonging to the clade I MPXV, which exhibits higher virulence in animal models of infection (7). Contrary to historical mpox cases, these new variants have gained an unprecedented ability to transmit amongst humans and controlling their spread is hampered by insufficient access to vaccines and efficacious antivirals (8). The emergence of MPXV variants that are resistant to the major antiviral tecovirimat is of particular concern and there is thus a great need for the discovery of novel antiviral strategies.

Innate immunity constitutes an important first line of defence against viral infection. A crucial family of innate immune cytokines are the interferons (IFNs), which are induced by key innate transcription factors and mediate potent antiviral activity, induce apoptosis in infected cells, and recruit inflammatory and antigen presenting cells to activate an adaptive immune response to clear the infection (9). Upon binding to cognate receptors, IFNs activate Janus kinase (JAK) and signal transducer and activator of transcription (STAT) proteins, resulting in the expression of hundreds of IFN-stimulated genes (ISGs), many of which are directly antiviral (10). One of the major host sensors for OPXV is the 2’3’ cyclic GMP-AMP (cGAMP) synthase (cGAS) (11–13), which upon binding double-stranded DNA, catalyses the production of second messenger cGAMP from ATP and GTP (14, 15). cGAMP then binds and activates stimulator of IFN genes (STING) in the ER, inducing its dimerisation, phosphorylation and translocation to the Golgi where IFN responsive factor (IRF)3 is recruited and subsequently phosphorylated (16, 17). Phosphorylated IRF3 then translocates to the nucleus and induces the transcription of IRF-dependent genes including the type I IFNs.

OPXV are characterised by their ability to antagonise and subvert host immunity via the deployment of an armoury of viral immunomodulators (18–21). Many OPXV, including vaccinia virus (VACV) and ectromelia virus (ECTV), express a viral poxin (OPG 188, *B2R* in VACV), a cGAMP-specific nuclease that cleaves the 3′–5′ bond and prevents recognition by STING (22, 23). In some OPXV, including MPXV and ECTV, poxin is fused to a domain with similarity to the mammalian Schlafen (Slfn) proteins, hence termed viral Slfn (vSlfn) (24–26). Deletion of *vSlfn* from ECTV results in a >5-log reduction in virulence in a lethal mouse model of infection, which maps to the poxin domain, demonstrating the importance of the cGAS-STING pathway in protection against OPXV infection (24). OPXV also express F17, which dysregulates mTOR simultaneously destabilising cGAS (27), and E5, which triggers cGAS ubiquitination and proteasome-dependent degradation (28). In addition, OPXV express multiple factors targeting IRF3 (29, 30) or able to supress IRF3 activation downstream of the nucleic acid sensors DNA-PK (31), RNA polymerase III (32) and IFI16 (33), all of which contribute to virulence in mouse models of infection.

Whilst wild-type OPXV induce very little IFN expression in cell culture, they remain partially sensitive to its antiviral effects, including MPXV (34, 35). Here we demonstrate that poxin is an Achilles’ heel in MPXV infection and that strategies by-passing poxin activity during MPXV infection induce an IFN response and are effective in abrogating viral replication. We therefore reveal STING agonism as a new potential avenue for the treatment of mpox.

## Results

### Depletion of poxin during MPXV infection restores STING and IRF3 activation

OPXV poxin is expressed in isolation (VACV B2) or fused to a Slfn-like domain, originally termed vSlfn (Fig. S1A). The vSlfn protein is conserved (> 90% aa identity) amongst MPXV strains including clade I (Supplementary Fig. S1B). We initially cloned the MPXV vSlfn gene and observed that ectopic expression significantly dampened IFNβ activation induced by cGAS/STING to similar levels as ECTV vSlfn and VACV poxin (Fig. 1A). ECTV vSlfn lacking the poxin domain (ECTV vSlfnΔp26, Supplementary Fig. S1A) was inactive in this assay as expected (24). To assess the role of vSlfn during MPXV infection, we used short hairpin (sh)RNA targeting the poxin sequence and employed a recombinant VACV expressing FLAG-tagged poxin (Methods) to validate the efficiency of the knock-down (Supplementary Fig. S2C). In contrast to infection in shCtrl cells, shPoxin cells restored phosphorylation of STING and IRF3 upon MPXV infection despite the presence of other viral antagonists, with the two MPXV clade IIb lineages tested showing equivalent results (Fig. 1B). Furthermore, both MPXV lineages were less able to suppress HT-DNA-induced STING phosphorylation in the absence of vSlfn (shPoxin cells) (Fig. 1C). These findings demonstrate that MPXV antagonises IRF3 activation using poxin.

**Fig 1:**
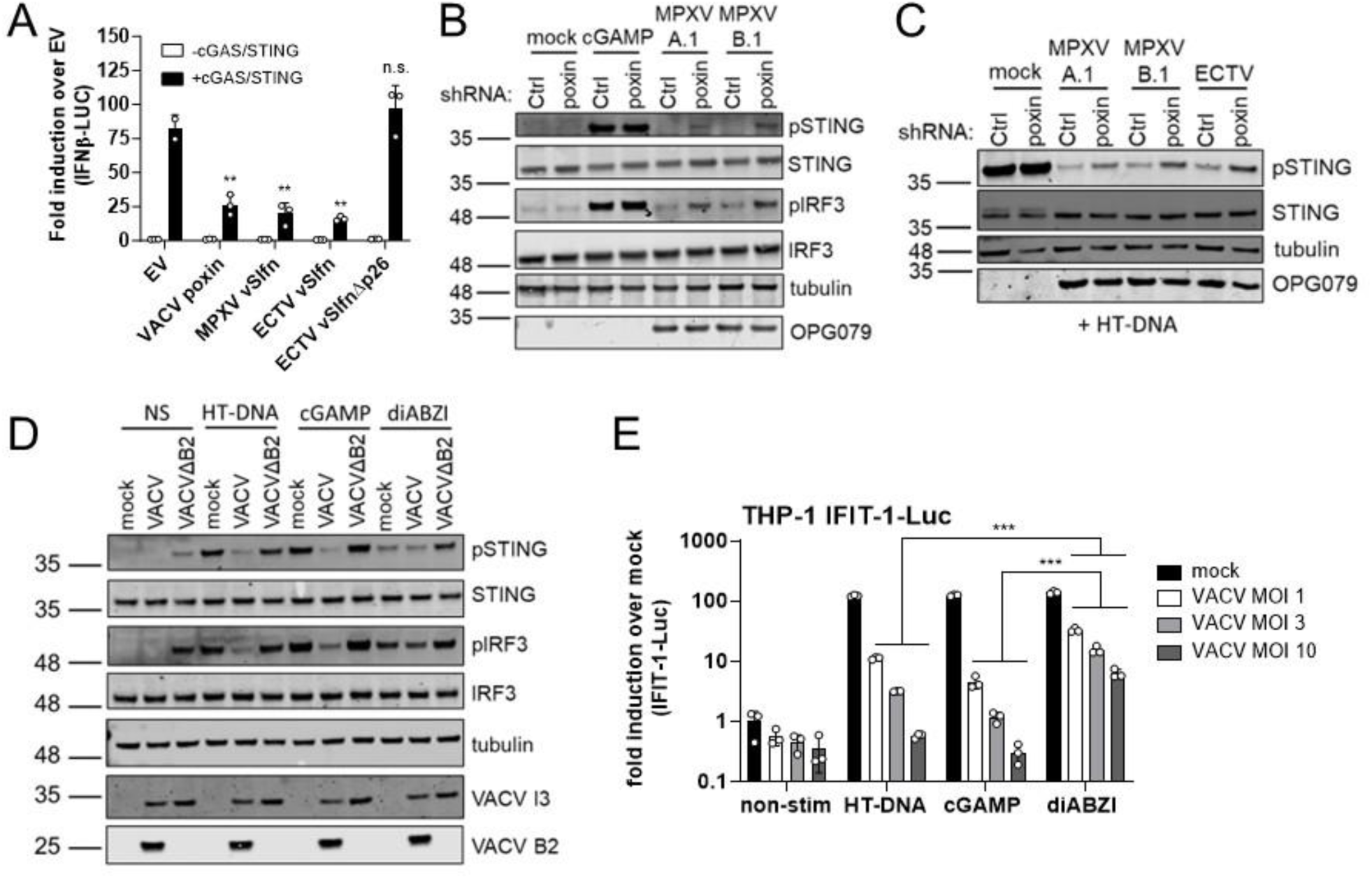
Non-CDN STING agonist diABZI bypasses poxin antagonism. (**A**) IFNβ reporter activity from HEK293T cells transfected for 24 h with 12.5 ng plasmid expressing VACV poxin, MPXV vSlfn, ECTV vSlfn or ECTV vSlfn lacking the p26 (poxin) domain (ECTV vSlfnΔp26) or an empty vector (EV) control per well and stimulated by co-transfection with 0.6 ng cGAS and 20 ng STING, or 20.6 ng EV as a non-stimulated control. (**B**) Immunoblot of THP-1-derived macrophages stably expressing either a control (Ctrl) or poxin-targeting shRNA, infected for 8 h at MOI 5 with the indicated viruses or stimulated for 2 h by transfection with 1 μg/ml cGAMP. Molecular mass markers are indicated on the left. (**C**) Immunoblot of THP-1-derived macrophages stably expressing either a control (Ctrl) or poxin-targeting shRNA, infected for 6 h at MOI 5 with the indicated viruses and then stimulated for 2 h by transfection with 1 μg/ml HT-DNA. Molecular mass markers are indicated on the left. (**D**) Immunoblot of THP-1-derived macrophages infected for 6 h with the indicated viruses at MOI 5 and then stimulated for 2 h with HT-DNA (transfected, 300 ng/ml), cGAMP (transfected, 2000 ng/ml) or diABZI (20 ng/ml), or left non-stimulated (NS). Molecular mass markers are indicated on the left. (**E**) IFIT-1 luciferase reporter activity from THP-1-derived macrophages infected for 8 h with VACV at the indicated MOIs and then stimulated for 16 h with HT-DNA (transfected, 300 ng/ml), cGAMP (transfected, 1000 ng/ml) or diABZI (50 ng/ml), or left non-stimulated (non-stim). Data are mean±SD, n = 3. Statistical analyses were performed using an unpaired Student’s t-test comparing each viral protein to the EV (**A**) or a one-way ANOVA with multiple comparisons (**E**). \*\**P* < 0.01, \*\*\**P* < 0.001, n.s. non-significant. All data presented are representative of at least 3 experimental repeats.

### Non-CDN STING agonist diABZI by-passes poxin antagonism

The development of compounds that activate STING is the focus of intense research. diABZI is an amidobenzimidazole derivative with enhanced binding to STING that elicits strong anti-tumour activity (36) and potently inhibits SARS-CoV-2 infection in respiratory epithelial cells and mice with very low toxicity (37, 38). Importantly, diABZI is not a cyclic compound like the canonical 2’3’-cGAMP and should be resistant to vSlfn activity, which we tested by comparing the ability of VACV and VACVΔB2 to suppress diABZI-mediated IRF3 activation. As expected VACV infection did not result in STING or IRF3 phosphorylation in non-stimulated cells and significantly reduced phosphorylation induced by HT-DNA and cGAMP (Fig. 1D). Conversely, VACV did not affect activation by diABZI. VACVΔB2 was unable to inhibit STING or IRF3 activation by any of the agonists and even induced their phosphorylation in non-stimulated cells (Fig. 1D). The ability of diABZI to by-pass VACV antagonism impacted IRF3 transcriptional activity, because diABZI induced comparatively higher activation of a luciferase reporter under the control of the IFN-induced with tetratricopeptide repeats 1 (*IFIT1*) promoter (13, 39) relative to equivalent doses of HT-DNA and cGAMP in cells infected with VACV at different multiplicities (Fig. 1E). In conclusion, a non-CDN STING agonist activated IRF3 responses despite the presence of poxin during MPXV infection.

### Non-CDN STING agonist activity is antiviral against OPXV

OPXV infection typically initiates in fibroblasts and keratinocytes at the lesion site and disseminates systemically via monocytes and macrophages reaching distal sites including secondary target organs. diABZI showed no toxicity in THP-1 derived macrophages (Fig. 2A) or primary human foetal foreskin fibroblasts (HFFF, Fig. 2B) at a range of doses that induced a potent innate immune response as measured by *IFIT1*-Luc (Fig. 2C) and endogenous ISG expression (Supplementary Fig. S2A and S2D). We therefore assessed the antiviral activity of diABZI using a recombinant VACV carrying firefly luciferase under the control of the viral thymidine kinase (*TK*) promoter. In both THPs and HFFF, diABZI potently suppressed luciferase activity after low multiplicity of infection (MOI), mimicking spreading infection (Fig. 2D and E). diABZI was also effective at suppressing higher MOI, single round infection (Fig. 2F and G). Inhibition of viral gene expression was confirmed measuring both early and late viral transcripts (Fig. 2H). Remarkably, when diABZI antiviral activity was determined measuring infectious progeny titre, this was shown to completely abrogate VACV replication, both at high (Fig. 2I) and low (Fig. 2J) multiplicity. diABZI was most effective following pre-treatment of the cells but retained significant restrictive capacity post-infection (Fig. 2J).

**Fig 2:**
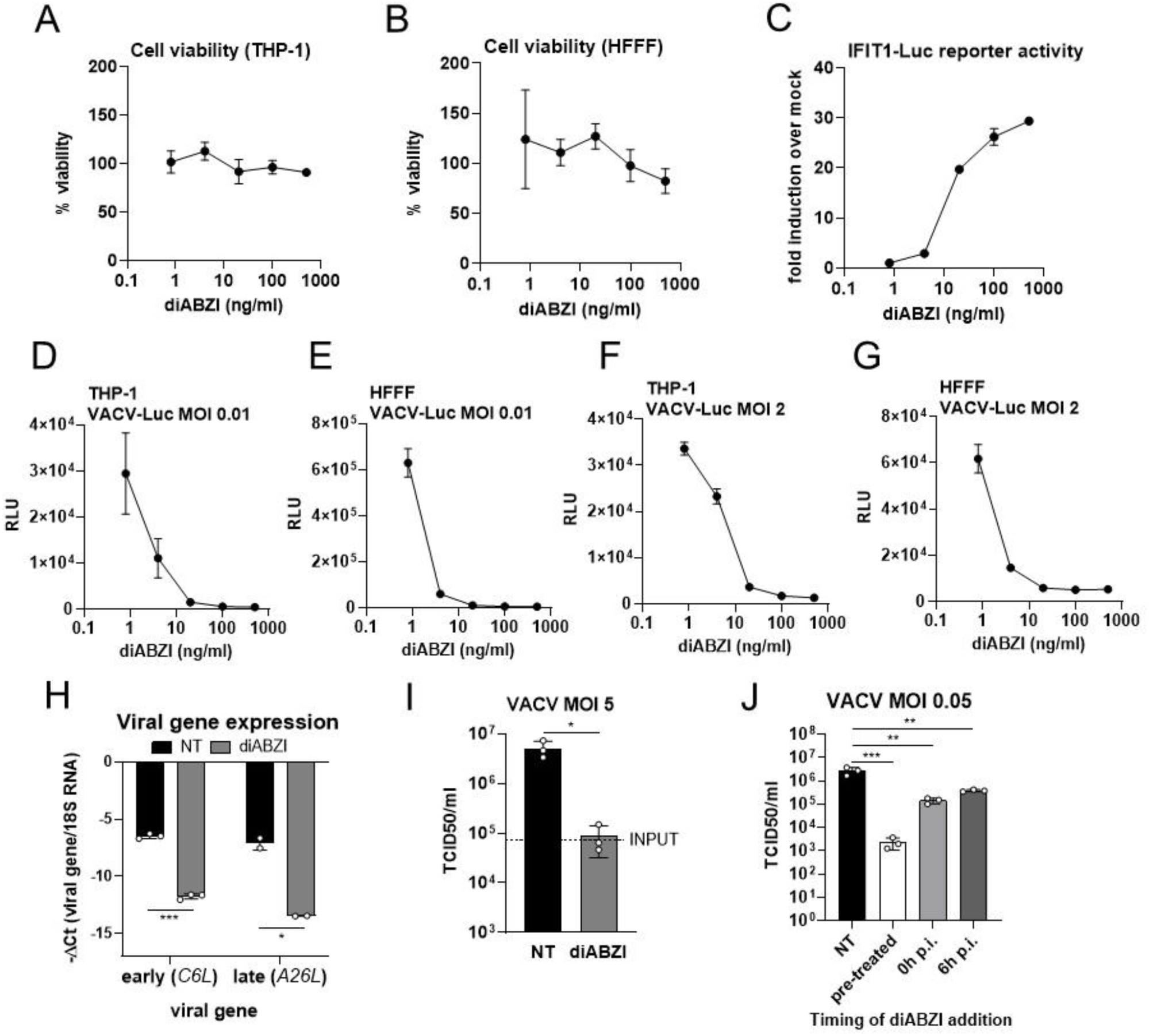
diABZI is antiviral against OPXV. (**A,B**) Cell viability at 48 h of THP-1-derived macrophages (**A**) and primary fibroblasts (HFFF, **B**) treated with the indicated doses of diABZI. (**C**) IFIT-1 luciferase reporter activity from THP-1-derived macrophages treated for 16 h with the indicated doses of diABZI. (*D,E*) Firefly luciferase activity (relative light units, RLU) from THP-1-derived macrophages (**D**) or HFFF (**E**) pre-treated for 16 h with the indicated doses of diABZI and then infected with VACV-Luc for 24 h at an MOI of 0.01. (**F,G**) Firefly luciferase activity (relative light units, RLU) from THP-1-derived macrophages (**F**) or HFFF (**G**) pre-treated for 16 h with the indicated doses of diABZI and then infected with VACV-Luc for 8 h at an MOI of 2. (**H**) Viral gene expression measured by RT-qPCR from THP-1-derived macrophages pre-treated for 16 h with 100 ng/ml diABZI or left non-treated (NT) and then infected with VACV at MOI 5 for either 2 hpi (early viral transcript *C6L*) or 6 hpi (late viral transcript *A26L*). (**I**) Viral titres at 24 hpi from THP-1-derived macrophages pre-treated for 16 h with 100 ng/ml diABZI or left non-treated (NT) and then infected with VACV at MOI 5. Input titre at 0 hpi is indicated with a dotted line. (**J**) Viral titre at 48 h in THP-1-derived macrophages infected with VACV at MOI 0.05 and either not treated (NT) or treated with 100 ng/ml diABZI either for 16 h prior to infection (pre-treated), at the time of infection (0h p.i.) or 6 h post-infection (6h p.i.). Data are mean±SD, n = 3 (except *A26L* qPCR where n=2). Statistical analyses were performed using an unpaired Student’s *t*-test (**H,I**) or a one-way ANOVA with multiple comparisons (**J**). \**P* < 0.05, \*\**P* < 0.01 *** *P* < 0.001. All data presented are representative of at least 3 experimental repeats.

### diABZI and IFNβ induce overlapping and unique mRNA expression profiles

To obtain a global view of diABZI responses in both cell types, we performed total mRNA sequencing and compared this to the type I IFN, IFNβ. Quantitation of representative ISG induction in THP-1 cells (Supplementary Fig. S2A-B) and HFFF (Supplementary Fig. S2D-E) demonstrated that responses were maximal at a dose of 100 ng/ml of diABZI and 5 ng/ml of IFNβ, and were largely similar in magnitude between diABZI and IFNβ in both cell types (Supplementary Fig. 2SC and S2F). These doses were thus selected for comparative RNA sequencing studies. diABZI induced large transcriptional changes, with the upregulation of >1,800 and >1,400 genes in differentiated THP-1 cells and fibroblasts, respectively (Fig. 3A and B). This was greater than the number of genes induced in either cell type by IFNβ (>1,600 genes in differentiated THP-1 cells and >600 in fibroblasts, Supplementary Fig. S3A-B, S3D-E). Interestingly, the overlap of diABZI-and IFNβ-induced genes was greater in differentiated THP-1 cells than fibroblasts, with >900 unique genes induced by diABZI in HFFF (Fig. 3C). Analysis of the top 20 upregulated KEGG pathways revealed strong activation of various immune, cytokine and pathogen-related pathways, as expected (Fig. 3D, E), showing significant overlap with the top upregulated pathways induced by IFNβ (Supplementary Fig. S3C, S3F). In addition to the convergent activation of canonical ISGs such as the IFITs, TRIM5, CXCL10 and BST2 (tetherin), diABZI induced the expression of IFNs β and λ1, and a plethora of cytokines including members of the CCL, CXCL and interleukin families, particularly in fibroblasts (Fig. 3F). Many of these cytokines were uniquely induced in HFFF by diABZI and were not observed following IFNβ treatment.

**Fig 3:**
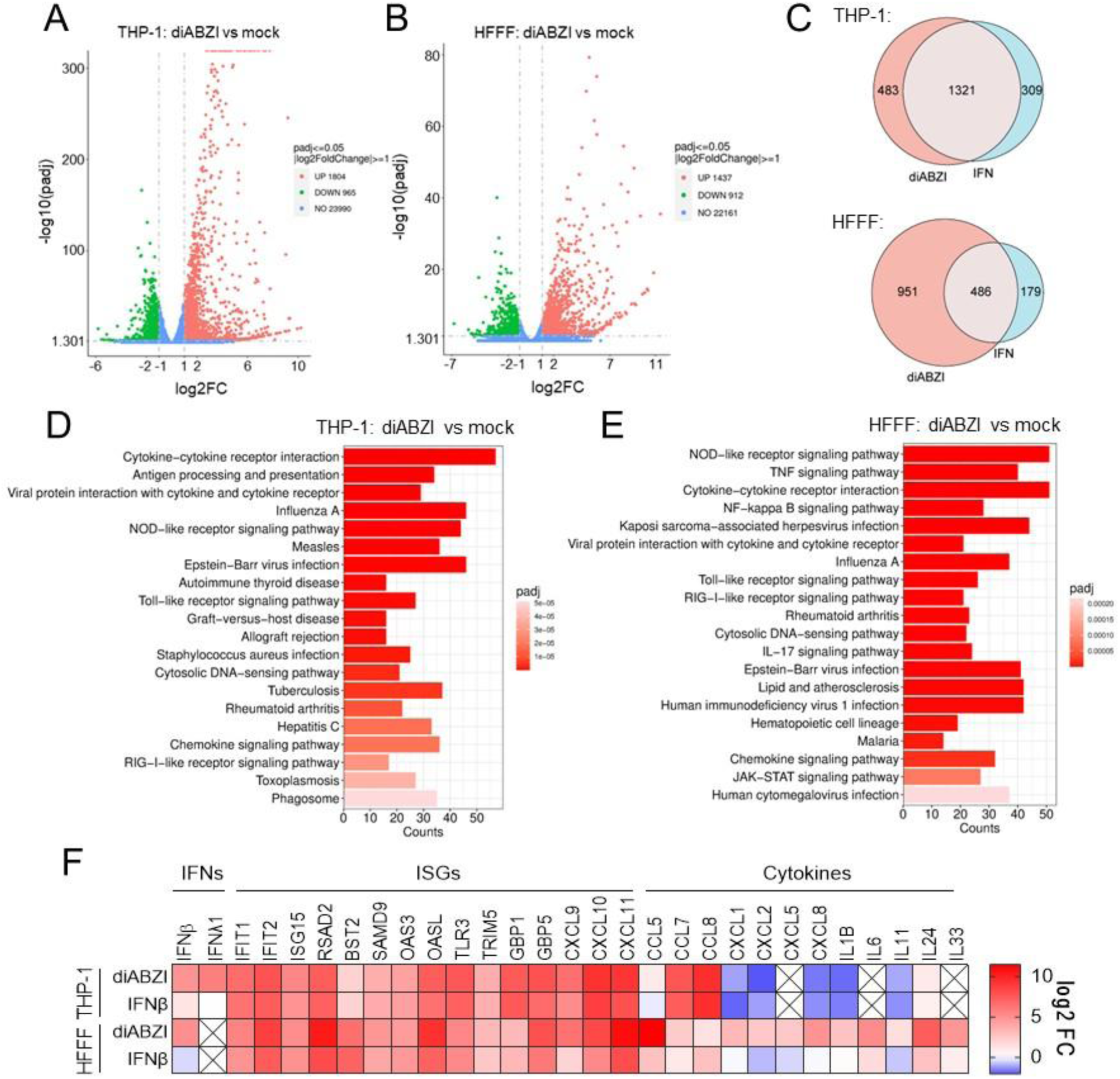
diABZI and IFNβ induce overlapping and distinct mRNA expression profiles. (**A,B**) Significantly differentially expressed genes (log2FC>1) by volcano plot analysis at 16 h post-stimulation with 100 ng/ml diABZI in THP-1-derived macrophages (**A**) and HFFF (**B**) compared with mock cells. *P* value determined by DESeq2. (**C**) Venn diagram of significantly upregulated genes (log2FC>1) at 16 h post-stimulation with 100 ng/ml diABZI or 5 ng/ml IFNβ in THP-1-derived macrophages and HFFF. (**D,E**) Top 20 significantly upregulated pathways (KEGG analysis) from RNAseq data at 16 h post-stimulation with 100 ng/ml diABZI in THP-1-derived macrophages (**C**) and HFFF (**D**) compared with mock cells. *P* value determined by DESeq2. (**F**) Heat map showing log2 fold change (FC) of selected genes from RNAseq data of THP-1-derived macrophages and HFFF cells stimulated with 100 ng/ml diABZI or 5 ng/ml IFNβ for 16 h. Boxes with an X indicate transcript not detected.

### Non-CDN STING agonist restriction of OPXV is IFN-dependent

We then comparatively determined the restrictive capacity of diABZI and IFNβ in cell culture. At the range of doses tested, both diABZI and IFNβ significantly restricted VACV infection in differentiated THP-1 cells (Fig. 4A) and fibroblasts (Fig. 4B). Restriction in macrophages was always between 1-2 log higher than in fibroblasts, in line with the greater number of differentially expressed genes observed at transcriptional level in this cell type (Fig. 3). No statistically significant difference was observed between the two treatments, suggesting that diABZI restriction relied on the IFN pathway. To address this, we employed genetically-modified cells and pharmacological inhibitors. As expected diABZI restrictive capacity was completely impaired in THP-1 cells lacking STING and viral replication was restored to the same levels observed in mock-treated cells (Fig. 4C). A similar rescue of viral replication was observed in diABZI-treated THP-1 cells lacking IRF3 or IFNAR, or in cells treated with the JAK1 inhibitor ruxolitinib (Fig. 4D). diABZI restriction was also rescued to levels of the mock-treated cells by ruxolitinib in fibroblasts (Fig. 4E). These findings indicated that diABZI restriction of OPXV was type I IFN dependent in these cell types.

**Fig 4:**
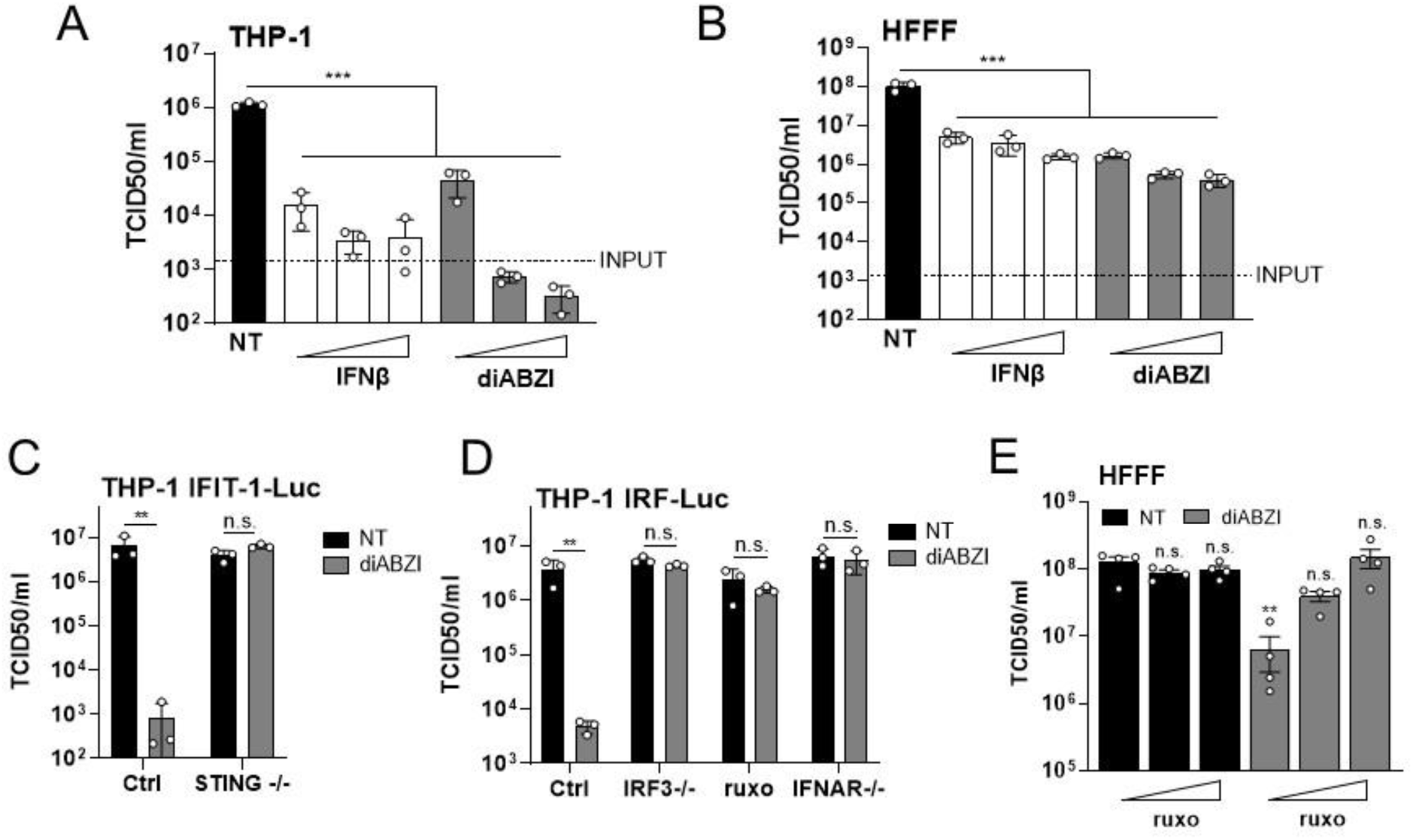
diABZI restriction of OPXV is type I IFN-dependent. (**A,B**) Viral titres at 48 h in THP-1-derived macrophages (**A**) or HFFF (**B**) either not treated (NT) or pre-treated for 16 h with diABZI (20, 100, 500 ng/ml) or IFNβ (1, 5, 25 ng/ml) and then infected with VACV at MOI 0.05. Input titre at 0 hpi is indicated with a dotted line. (**C,D**) Viral titres at 48 hpi from control (Ctrl), STING-/-(**C**), IRF3-/-, IFNAR-/- or ruxolitinib-treated (ruxo) (**D**) (2 μM, added at time of diABZI pre-treatment) THP-1-derived macrophages pre-treated for 16 h with 100 ng/ml diABZI or left non-treated (NT) and then infected with VACV at MOI 0.05. (**E**) Viral titres at 48 hpi in HFFF either not treated (NT) or pre-treated for 16 h with 100ng/ml diABZI-/+ 0, 2 or 5 μM ruxolitinib (ruxo) and then infected with VACV at MOI 0.05. All data are mean ± SD, n = 3 per condition. Statistical analyses were performed using a one-way (*A,B,E*) or two-way (*C,D*) ANOVA with multiple comparisons. \*\**P* < 0.01 \*\*\**P* < 0.001. All data presented are representative of at least 2 experimental repeats.

### Non-CDN STING agonist activity is antiviral against MPXV *in vitro* and *in vivo*

The robust induction of antiviral gene expression and restrictive activity against surrogate OPXV indicated that non-CDN STING agonists could be effective against MPXV. To test this, we infected both differentiated THP-1 cells and fibroblasts with MPXV and measured progeny virus titre in the presence or absence of diABZI. In line with previous results, diABZI completely abolished MPXV replication in THP-1 cells (Fig. 5A) and showed significant reduction in HFFF (Fig. 5B). Although rodents comprise the major reservoir for MPXV, common inbred strains of mice including BALB/c and C57BL/6 are resistant to MPXV. The wild mice derived strain CAST/EiJ, however, exhibits morbidity and is susceptible to MPXV-induced disease (40), particularly to the more virulent clade I MPXV strains (41). Using this model, we observed significant weight loss from day 4 post-infection in animals inoculated with 10^5^ plaque forming units (PFU) of MPXV Clade Ib, which were eventually euthanised (MPXV NT, Fig 5C, D). All mice survived however when pre-treated with 0.5 mg/kg diABZI 24 h prior to infection (MPXV diABZI (pre)) and showed significantly reduced weight loss, which was further reduced by diABZI treatment both pre-and post-inoculation (MPXV diABZI (pre+post)). Furthermore, diABZI was also found to be effective even when administered 24 h post-infection, significantly reducing weight loss and increasing survival to 60 % (MPXV diABZI (post)). These results demonstrate that STING activation through poxin-resistant agonists offers a therapeutic antiviral strategy against human mpox.

**Fig 5:**
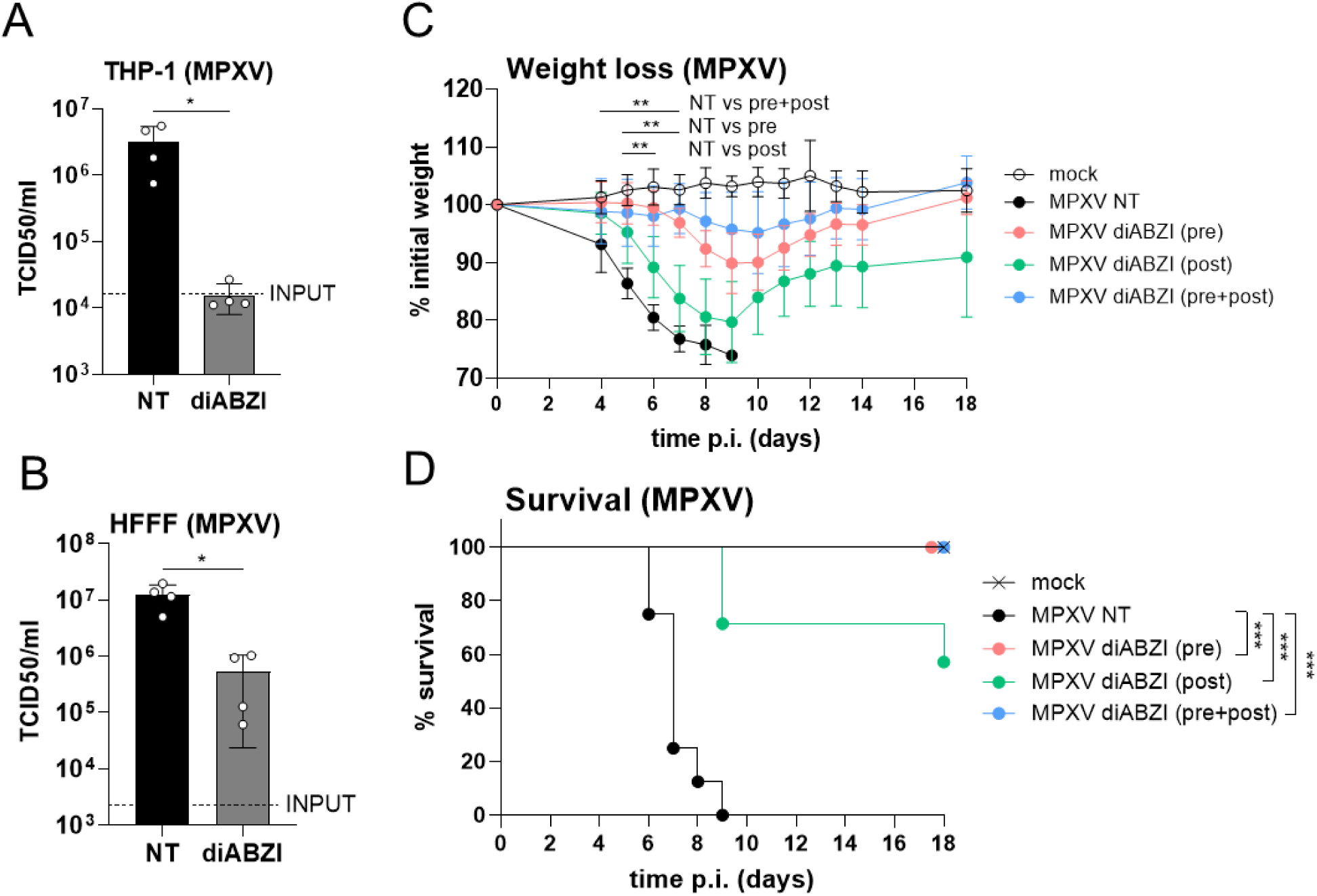
diABZI is antiviral against MPXV. (**A**, **B**) Viral titres at 48 h in THP-1-derived macrophages (**A**) or HFFF (**B**) either not treated (NT) or pre-treated for 16 h with diABZI (100 ng/ml) and then infected with MPXV (Clade IIb lineage B.1) at MOI 0.05. Input titre at 0 hpi is indicated with a dotted line. Data are mean ± SD, n = 4. Statistical analyses were performed using an unpaired Student’s t-test. *P < 0.05. Data are representative of 2 experimental repeats. (**C**, **D**) Groups of CAST mice (n=8) that had either been non-treated (NT) or treated for 24 h prior to infection (pre), 24 h post-infection (post) or both 24 h before and 24 h after infection (pre+post) with 0.5 mg/kg diABZI were infected intranasally with 10^5^ p.f.u. of MPXV (Clade Ib). Mice were monitored daily for bodyweight (**C**). Weight data are expressed as the mean ±SEM of the animals’ weights compared to their original weight at the day of inoculation. Statistical analyses were performed using Multiple Mann-Whitney tests. The horizontal line represents the days on which the indicated comparisons were statistically significant. **P < 0.01. Data are representative of 2 experimental repeats. (**D**), Survival data from (**C**). Statistical analyses were performed using a Mantel-Cox test. ***P < 0.001.

## Discussion

The innate immune system is a potent barrier to pathogen invasion. Pharmacological activation of innate immune pathways offers a potent strategy against viral infection that, in addition, by targeting the host rather than the virus, minimises the risk of developing antiviral resistance. STING is one of the major factors in the host innate response, which has led to the development of small-molecule agonists with higher potency and better bioavailability than its endogenous cognate activator cGAMP (36). Here we took advantage of the acyclic nature of the STING agonist diABZI to disable poxin, a major OPXV STING antagonist, and probe an Achilles’ heel in OPXV immune evasion. Our data highlight the remarkable restrictive capacity of this compound, particularly in monocyte-derived macrophages where MPXV replication was abolished. Although all OPXV zoonoses cause a local lesion at the infection site usually within 3 days after the onset of prodromal symptoms, a major clinical difference between mpox and other OPXVs is a subsequent disseminated infection resembling human smallpox (42). Whilst the mechanisms that mediate such dissemination in humans are unclear, migratory inflammatory monocytes recruited to infected keratinocytic lesions are permissive to OPXV infection (43), making them potential vehicles for long-distance in-host spread. Elimination of these mobile infected cells is a therapeutically attractive goal to curb spread and secondary lesion formation, and our data indicates that this STING agonism may be particularly effective at it.

STING activation was also restrictive in fibroblasts but did not completely suppress viral replication as in differentiated THP-1 cells. This observation correlated with a broader and more robust ISG induction in THP-1 cells. Several OPXV restriction factors have been reported including IFITs, SAMD9 and TRIM5α. These ISGs were equally induced by diABZI and IFNβ in both cell types and therefore other restrictive ISGs are likely to exist and contribute to the differing restrictive capacities of macrophages and fibroblasts observed in this study. Besides a canonical type I IFN response, diABZI induced signalling pathways (e.g. NF-κB) that although not directly antiviral in cells, might be important in shaping the overall immune response and limiting toxicity. This was reflected in the upregulation of multiple cytokines with chemoattractant capacity that would promote leukocyte infiltration and viral clearance in vivo. In addition, STING activation is expected to benefit from IRF3-dependent but STAT1-independent induction of ISGs (44) potentially enabling antiviral activity in cells with poor IFN responsiveness.

Human mpox was traditionally a zoonotic disease from rural areas in West and Central Africa. Since smallpox eradication the incidence of mpox has progressively increased (45), culminating in several large multi-country outbreaks in recent years. These upsurges prompted the World Health Organization (WHO) to declare mpox a public health emergency of international concern twice in 2022 and 2024. At present, mpox outbreaks are contained through a combination of public health measures and vaccination. There are three WHO-approved vaccines: ACAM2000, JYNNEOS, and LC16m8. Each of these, however, presents limitations: ACAM2000 has significant adverse effects and was originally approved as a suitable replacement for Dryvax in the event of bioterror (46); JYNNEOS (also known as Imvanex) is not approved for children under 12 years old, requires 2 doses administered 4-weeks apart and is difficult to manufacture, limiting its global distribution and accessibility (47); and LC16m8 requires administration by scarification using a bifurcated needle and is not approved during pregnancy or in people who are immunocompromised (47). Complementing these vaccines, several therapeutic agents are approved against mpox. Tecovirimat (also known as ST-246 or TPOXX) is the most extensively used. It prevents virus egress by targeting the viral protein F13, which is essential for production of extracellular virus and in-host dissemination (48). Tecovirimat is safe, orally bioavailable and protects multiple animal species form lethal OPXV challenge (48), but its clinical efficacy in humans needs to be re-assessed after it failed to meet several clinical endpoints in two recent trials (49). Furthermore, concerns exist over its low barrier to viral resistance and the emergence and transmission of resistant MPXV strains (35, 50). These limitations and the current evolution of MPXV as a growing public health emergency press for the development of novel therapeutic agents. Our work provides evidence that strategies to disable viral antagonism of STING are effective antivirals and offer a promising new avenue for anti-mpox therapeutics in isolation or in synergistic combination.

## Acknowledgments

We thank D. Ulaeto (Defence Science and Technology Laboratory, United Kingdom), M. Weekes (University of Cambridge, United Kingdom) and G.L. Smith (Shanghai Institute for Immunity and Infection, China) for providing reagents, and K. Maringer (The Pirbright Institute, United Kingdom) for support with the BSL-3 facility. This work was funded by the UK BBSRC grants BB/X0011356/1 and BB/V015265/1 to CMdM; the UK MRC grant MR/Z503885/1 to CMdM and RPS; the Royal Society Research Grant RGS\R2\252220 to RPS; the Spanish Ministry of Science, Innovation and Universities grants PID2022-136867NB-I00 and RTI2021-128580OB-100 to BH and AA, respectively, and La Caixa Foundation grant HR23-00558 to AA. Work at the Pirbright Institute was supported by institutional BBSRC grants BBS/E/PI/23NB0004 and BBS/E/PI/23NB0003. For the purpose of open access, the author has applied a Creative Commons attribution license (CC BY) to any Author Accepted Manuscript version arising from this submission.

## Author Contributions

LE, RPS, BH and CMdM conceived the study. LE, ARU, AJMH, TSF, BH and RPS performed the experiments and analysed the data. CMdM, BH and RPS wrote the manuscript (original draft). AA, BH, CMdM and RPS reviewed and edited the paper (final version). CMdM, RPS, AA and BH acquired funding. All authors read and agreed with the paper.

## Methods

### Cells and reagents

Human foetal foreskin fibroblasts (HFFF-2), HEK 293T cells and BSC-40 cells were maintained in DMEM (Gibco) supplemented with 10 % foetal bovine serum (FBS, Labtech) and 100 U/ml penicillin plus 100 μg/ml streptomycin (Pen/Strep; Gibco). THP-1 cells were maintained in RPMI (Gibco) supplemented with 10 % FBS and Pen/Strep and differentiated for 48 h into macrophage-like cells in the presence of 50 ng/ml phorbol 12-myristate 13-acetate (PMA, Peprotech). THP-1-IFIT-1 cells (39) and THP-1-IFIT-1 STING-/-cells (51) have been previously described. THP-1 Dual Control, IFNAR-/-and IRF3-/-cells were obtained from Invivogen. JAK inhibitor ruxolitinib was from CELL guidance systems. IFNβ was obtained from Peprotech. Herring-testis DNA was obtained from Sigma. cGAMP and diABZI were from Invivogen. For stimulation of cells by transfection, transfection mixes were prepared using lipofectamine 2000 according to the manufacturer’s instructions (Invitrogen).

### Virus stocks

MPXV Clade IIb lineage B.1 (MPXV_CVR_S1; ON808413) was isolated in 2022 from a patient at the University of Glasgow-Centre for Virus Research. MPXV Clade IIb lineage A.1 (MPXV_UK_P3; MT903345) was isolated in 2018 at the UK Health Security Agency. MPXV Clade Ib (MpxV/PHAS-506/Passage-03/SWE/2024_09_11; Gisaid EPI_ISL_19348512) was previously described (52) and was sourced from the European virus archive global (EVAg). Other viruses used were VACV Western Reserve (WR, accession number AY243312), VACVΔB2 (12) and VACV-Luc (obtained from Geoffrey L Smith). For construction of VACV encoding Flag-tagged B2 from its natural locus (VACV.Flag.B2), a transient dominant selection strategy was used. First, a plasmid containing the coding sequence of *B2R* fused to 3 repeats of the Flag-tag at the N terminus, flanked by 200-bp of the 5’ and 3’ regions of the *B2R* locus, was cloned in a pUC13 vector containing the Escherichia coli guanylphosphoribosyl transferase (Ecogpt) gene fused in-frame with the enhanced green fluorescent protein (EGFP). Recombination into VACVΔB2, isolation and titration were then performed as detailed previously (53). Purification was performed by ultracentrifugation of cytoplasmic extracts over a 36 % (w/v) sucrose cushion (32,900 × g for 80 mins at 4 °C) prior to use. Infection experiments with MPXV were performed in a Biosafety level 3 laboratory at The Pirbright Institute, Surrey.

### Viral replication assays

For viral replication assays cells were seeded at 2.5×10^5^ cells/well in 24 well plates. The following day cells were treated overnight with diABZI, IFNβ, or mock treated with medium as indicated in the figure legends and then infected the next morning on ice at the indicated MOI in 200 μl/well medium. After 60 min the inoculum was removed and the cells washed once with medium to remove unbound virus before 500 μl medium was added per well and the cells transferred to 37 °C. IFNβ and diABZI were not replaced in this medium. Virus was harvested at 0 and 48 hpi by scraping the cells in their medium and freeze-thawing three times before calculation of viral titres by tissue culture infectious dose 50 (TCID50) assay according to the Reed and Muench method as previously described (54). TCID50 assays were performed on BSC-40 cells and incubated for 3 days for VACV and 6-7 days for MPXV. All infections were performed in medium supplemented with 2.5 % FBS. For experiments with ruxolitinib (either 2 or 5 μM as indicated in legends), this was added 1 h prior to diABZI pre-treatment and replaced after the initial infection. diABZI was not replaced after initial infection.

### Cell viability assay

Cell viability in the presence of diABZI was calculated using the CyQUANT™ MTT Cell Proliferation Assay Kit (Invitrogen) according to the manufacturer’s protocol.

### shRNA lentivirus transduction

Hairpin sequences against the VACV WR poxin gene (top: GAAGGAGTAGGGATTCATCAT; bottom: CAAAGAGAAGGCCAAAGAAAT) were designed using siRNA Wizard v3.1 Software (InvivoGen) and annealed into oligo duplexes. The duplexes were then cloned into an HIV-1-based shRNA expression vector encoding puromycin resistance (pSIREN, Clontech) at BamHI-EcoRI sites, and the products were checked by sequencing. VSV-G-pseudotyped lentiviral particles and transduced cell lines were obtained as previously described (12).

### Transcriptomics and bioinformatics

For transcriptomics experiments cells were seeded at 2.5×10^5^ cells/well in 24 well plates and stimulated in triplicate with 100 ng/ml diABZI or 5 ng/ml IFNβ for 16 h. RNA was purified using the RNeasy mini RNA extraction kit with on-column DNase treatment and sent for bulk RNA sequencing by Novogene. Messenger RNA was purified from total RNA using poly-T oligo-attached magnetic beads. After fragmentation, the first strand cDNA was synthesised using random hexamer primers, followed by the second strand cDNA synthesis using either dUTP for directional library or dTTP for non-directional library. The library was checked with Qubit and real-time PCR for quantification and bioanalyzer for size distribution detection. Quantified libraries were then pooled and sequenced on an Illumina platform. RNA-seq reads were aligned to the *Homo Sapiens* genome using HISAT2(55) and featureCounts v1.5.0-p3 was used to count the read numbers mapped to each gene and the FPKM of each gene was calculated based on the length of the gene and reads count mapped to it. Differentially expressed genes (DEG) were calculated using DESeq2(56). The resulting *P*-values from DESeq2 were adjusted using the Benjamini and Hochberg’s approach for controlling the false discovery rate. Genes with an adjusted *P*-value<0.05 and log2FC>1 were found by DESeq2 were assigned as differentially expressed. DEGs were then used for KEGG pathway enrichment analysis with *p*-values <0.05 considered as significant (57). KEGG pathway analysis figures were generated using SRPlot (accessed from https://www.bioinformatics.com.cn/srplot). Heat maps were generated using GraphPad Prism (version 10).

### Luciferase reporter assays

Dual luciferase reporter gene assays in HEK293T cells were performed as we have described previously (24). Plasmids encoding codon-optimised ECTV vSlfn and ECTV vSlfn lacking the poxin/p26 domain (ECTV vSlfnΔp26) have also been previously described (24). Codon-optimised VACV *B2R* (GeneArt, Invitrogen) and MPXV vSlfn (obtained from Mike Weekes) were cloned into a modified pcDNA4 vector with 3 copies of the FLAG tag at the N terminus (58). Assays involving THP-1-IFIT1-Luc or THP-1-Dual cells were performed in 96-well assay plates and luciferase activity was measured with coelenterazine (Nanolight Technologies, 2 μg/ml) in a CLARIOstar plate reader (BMG Labtech). For assays involving VACV-Luc, cells were first lysed with 100 μl/well passive lysis buffer (Promega) and freeze-thawed prior to luciferase readings. Luciferase activity was then measured with home-made firefly luciferase substrate (20 mM Tricine, 2.67 mM MgSO4.7H_2_O, 0.1 mM EDTA, 33.3 mM DTT, 530 μM ATP, 270 μM acetyl coenzyme A, 132 μg/ml Luciferin (Prolume Ltd.), 5 mM NaOH, 0.26 mM (MgCO3)4Mg(OH)2.5H_2_O)). Fold inductions were calculated by normalising to a mock-treated control.

### RT-qPCR

RNA was extracted from cells using a total RNA purification kit (QIAGEN) according to the manufacturer’s protocol. Five hundred ng RNA was used to synthesise cDNA using Superscript III reverse transcriptase (Invitrogen), also according to the manufacturer’s protocol. For viral gene expression qPCR, RNA was treated with DNase prior to cDNA synthesis (Invitrogen). cDNA was diluted 1:5 in water and 1.5 μl used as a template for real-time PCR using SYBR® Green PCR master mix (Applied Biosystems) and a Quant Studio 5 (Applied Biosystems) real-time PCR machine. Expression of each gene was normalised to an internal control (*GAPDH* or *18S-RNA*) and these values were then normalised to mock cells to yield a fold induction. The following primers were used:

*GAPDH* Fwd 5’-GGGAAACTGTGGCGTGAT-3’, Rev 5’-GGAGGAGTGGGTGTCGCTGTT-3’ *18S RNA*: Fwd 5’-GTAACCCGTTGAACCCCA-3’, Rev 5’-CCATCCAATCGGTAGTAGCG-3’ *CXCL-10* Fwd 5’-TGGCATTCAAGGAGTACCTC-3’, Rev 5’-TGTAGCAATGATCTCAACACG-3’ *MxA* Fwd 5’-ATCCTGGGATTTTGGGGCTT-3’, Rev 5’-CCGCTTGTCGCTGGTGTCG-3’ *IFITM3* Fwd 5’-ACCATGAATCACACTGTCCAAACCTT-3’, Rev 5’-CCAGCACAGCCACCTCG-3’ *A26L (OPG153)*: Fwd 5’-ATCTCCCATGTGGTGGAATAC-3’, Rev 5’-GTTGATAGGTTAGAACATCAC-3’ *C6L (OPG029)*: Fwd 5’-TGTATTCTACGATAGAGTGC-3’, Rev 5’-AGATAAACTCACTGTTTATGG-3’

### Immunoblotting

SDS-PAGE and immunoblotting were performed as previously described (13) using the following primary antibodies: mouse-anti-tubulin (EMD Millipore), rabbit-anti-STING (Cell Signaling), rabbit-anti-pSTING (Ser 366, Cell Signaling), rabbit-anti-IRF3 (Abcam), rabbit-anti-pIRF3 (Ser 386, Abcam), mouse-anti-FLAG M2 (Sigma), rabbit-anti VACV B2 (OPG 188) polyclonal serum (12) and VACV I3 (OPG079) serum (obtained from David Ulaeto). Primary antibodies were detected with goat-anti-mouse/rabbit IRdye 680/800 infrared dye secondary antibodies and membranes imaged using an Odyssey Infrared Imager (LI-COR Biosciences).

### MPXV infection of mice

The experiments described and Cast/EiJ mice breeding were carried out at the Centro de Biologia Molecular Severo Ochoa (CBM) in Madrid (Spain) and were approved by the Ethical Review Board of CBM and CSIC under reference PROEX 241.1/21. Groups of 7-12 weeks (12-16 g) male Cast/EiJ mice were intranasally (i.n.) infected with a single 20 μl dose containing 10^5^ PFU of MPXV Clade Ib. After infection, viral inocula used were back-titrated by plaque assay to verify the administered viral dose. Mice were housed in ventilated racks under biological safety level 3 containment facilities and monitored daily for weight. diABZI-4 (0.5 mg/kg, MedChemExpress) was administered i.n. to animals either 24 h before MPXV infection (pre-treatment), 24 h post-infection (post-treatment) or both 24 h before and 24 h after infection (pre-post treatment).

## Statistical analyses

Statistical analyses were performed using an unpaired Student’s t-test (with Welch’s correction where variances were unequal), a one-or two-way ANOVA with multiple comparisons, Multiple Mann-Whitney tests, or a Mantel-Cox test as indicated in figure legends using GraphPad Prism software. * *P* < 0.05, ** *P* < 0.01, *** *P* < 0.001, n.s. non-significant.

---

**Fig S1:**
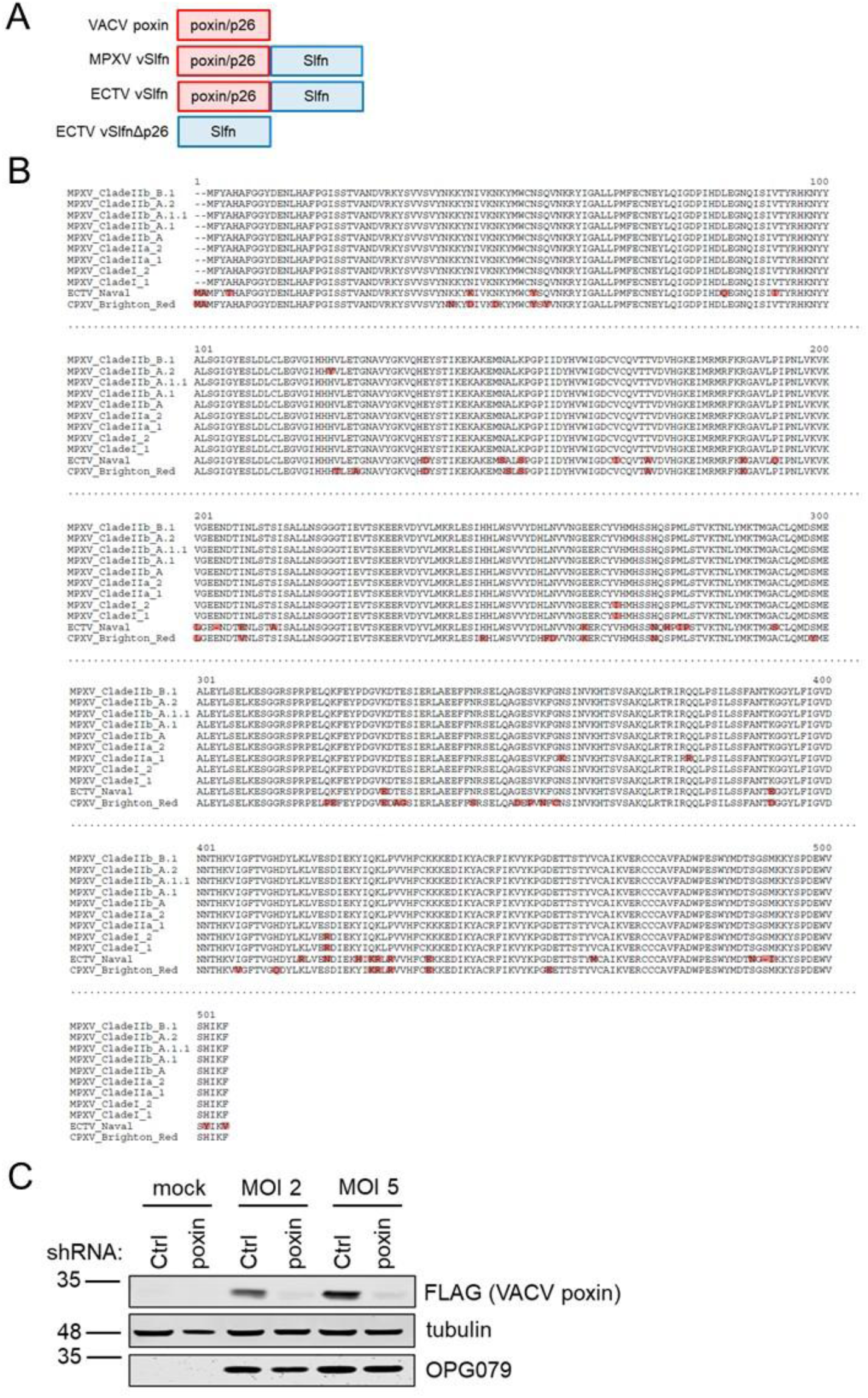
Non-CDN STING agonist diABZI bypasses poxin antagonism. (**A**) Schematic of the poxin/vSlfn constructs used in this study. (**B**) Amino acid alignment of OPXV vSlfn proteins with mismatches highlighted in red. (**C**) Immunoblot of THP-1-derived macrophages stably expressing either a control (Ctrl) or poxin-targeting shRNA, infected for 16 h with a VACV expressing FLAG-tagged poxin at MOI 2 or 5.

**Fig S2:**
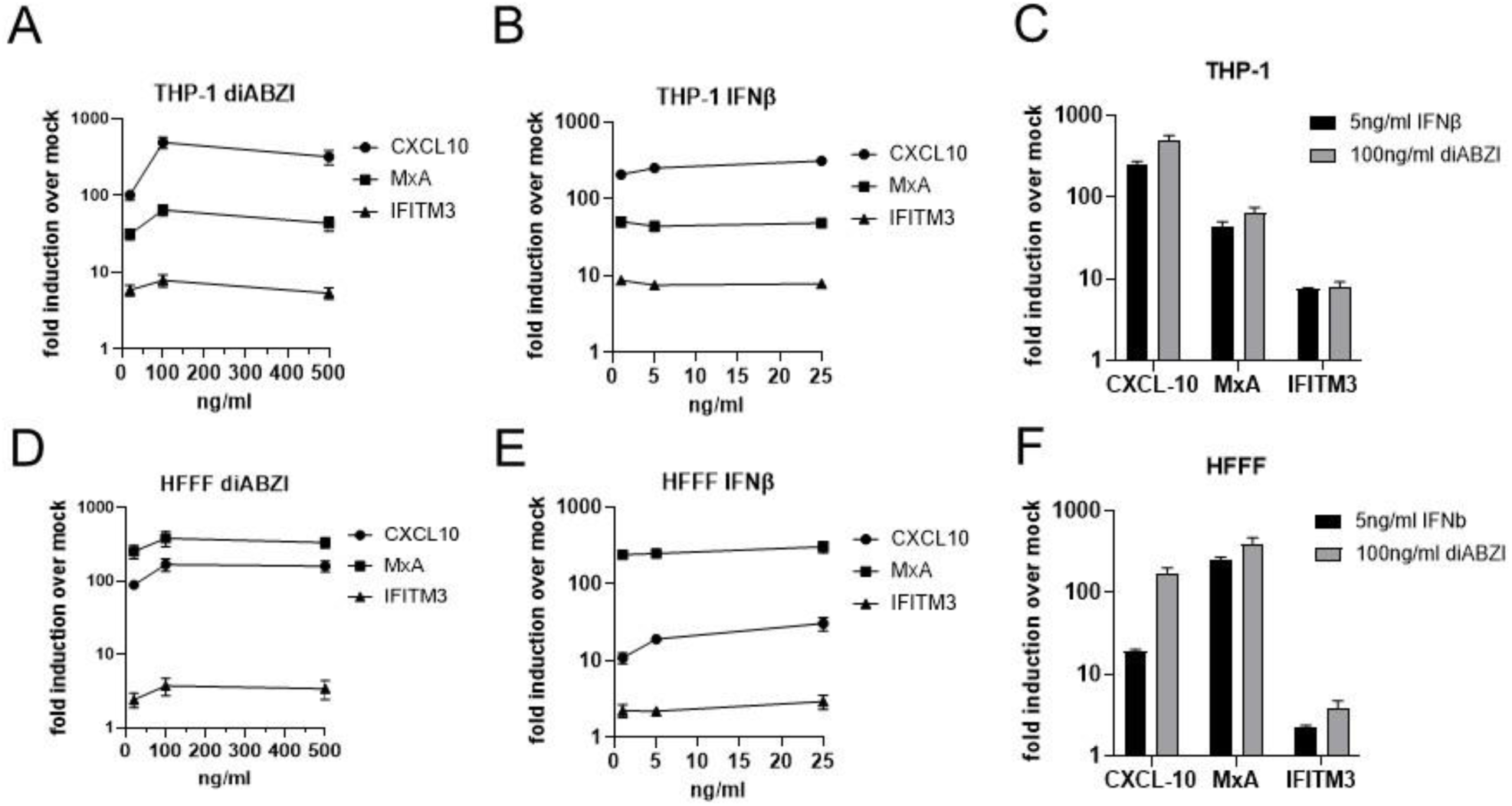
ISG induction by diABZI and IFNβ in THP-1 and HFFF. (**A,B**) ISG qPCR from THP-1-derived macrophages stimulated for 16 h with 20, 100 or 500 ng/ml diABZI (**A**) or 1, 5 or 25 ng/ml IFNβ (**B**). (**C**) Comparison of ISGs responses from (**A**) and (**B**) at the doses of diABZI and IFNβ selected for RNAseq. (**D,E**) ISG qPCR from HFFF stimulated for 16 h with 20, 100 or 500 ng/ml diABZI (**D**) or 1, 5 or 25 ng/ml IFNβ (**E**). (**F**) Comparison of ISGs responses from (*D*) and (*E*) at the doses of diABZI and IFNβ selected for RNAseq. Data are mean ± SD, n = 3. Data presented are representative of 2 experimental repeats.

**Fig S3:**
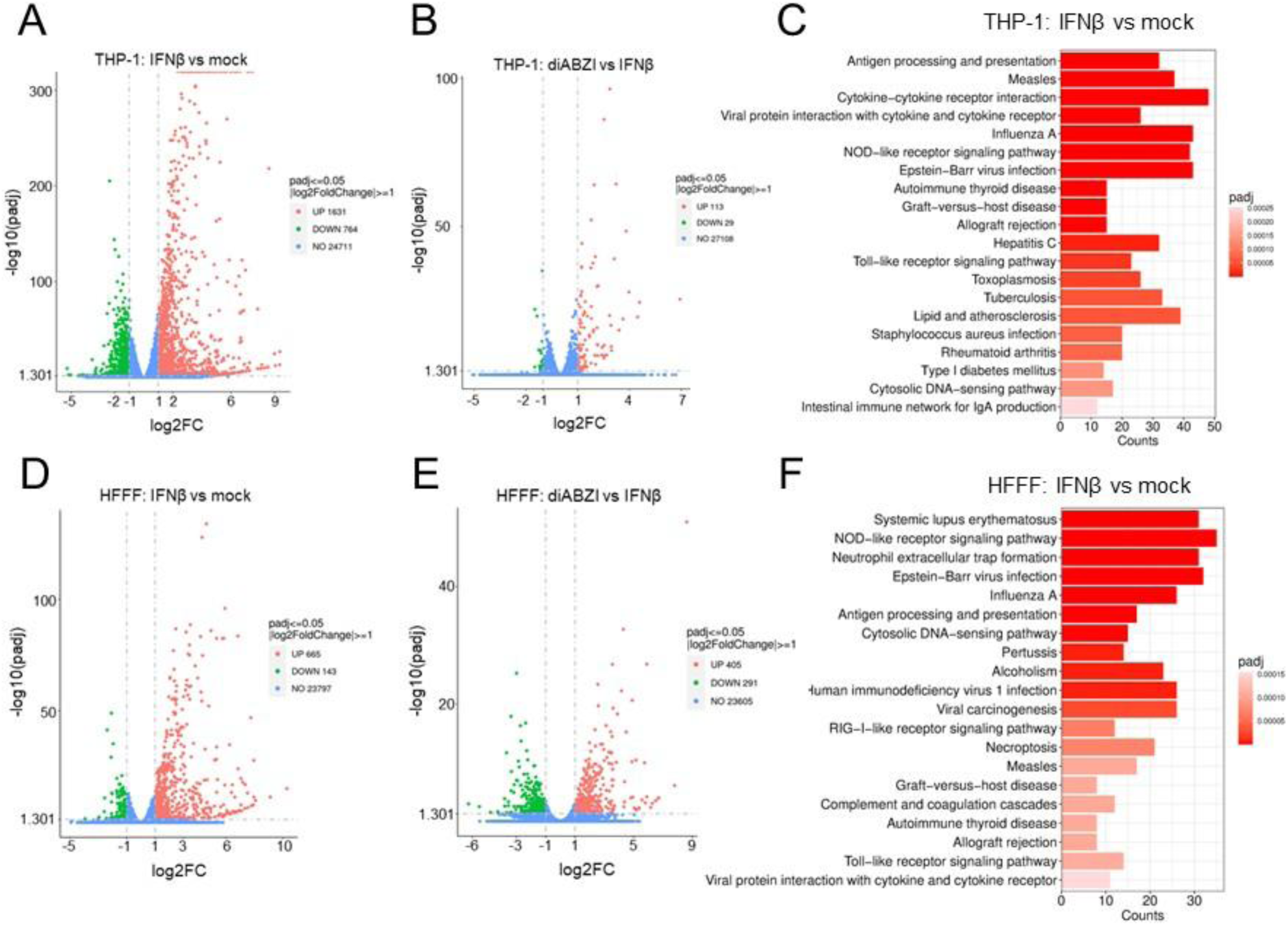
diABZI and IFNβ induce overlapping and distinct mRNA expression profiles. (**A,B**) Significantly differentially expressed genes (log2FC>1) by volcano plot analysis in THP-1-derived macrophages 16 h post-stimulation with 5 ng/ml IFNβ versus mock (**A**) or 100 ng/ml diABZI versus 5 ng/ml IFNβ (**B**). *P* value determined by DESeq2. (**C**) Top 20 significantly upregulated pathways (KEGG analysis) from RNAseq data at 16 h post-stimulation with 5 ng/ml IFNβ in THP-1-derived macrophages compared with mock cells. *P* value determined by DESeq2. (**D,E**) Significantly differentially expressed genes (log2FC>1) by volcano plot analysis in HFFF 16 h post-stimulation with 5 ng/ml IFNβ versus mock (**D**) or 100 ng/ml diABZI versus 5 ng/ml IFNβ (**E**). *P* value determined by DESeq2. (**F**) Top 20 significantly upregulated pathways (KEGG analysis) from RNAseq data at 16 h post-stimulation with 5 ng/ml IFNβ in HFFF compared with mock cells. *P* value determined by DESeq2.

